# Expansion of human centromeric arrays in cells undergoing break-induced replication

**DOI:** 10.1101/2023.11.11.566714

**Authors:** Soyeon Showman, Paul B. Talbert, Yiling Xu, Richard O. Adeyemi, Steven Henikoff

## Abstract

Human centromeres are located within α-satellite arrays and evolve rapidly, which can lead to individual variation in array lengths. Proposed mechanisms for such alterations in lengths are unequal cross-over between sister chromatids, gene conversion, and break-induced replication. However, the underlying molecular mechanisms responsible for the massive, complex, and homogeneous organization of centromeric arrays have not been experimentally validated. Here, we use droplet digital PCR assays to demonstrate that centromeric arrays can expand and contract within ~20 somatic cell divisions of a cell line. We find that the frequency of array variation among single-cell-derived subclones ranges from a minimum of ~7% to a maximum of ~100%. Further clonal evolution revealed that centromere expansion is favored over contraction. We find that the homologous recombination protein RAD52 and the helicase PIF1 are required for extensive array change, suggesting that centromere sequence evolution can occur via break-induced replication.

## Introduction

Centromeres are chromosomal regions where kinetochore assembly and microtubule attachments occur to ensure faithful genetic transmission of chromosomes to daughter cells during mitosis and meiosis^1^. Active centromeres are epigenetically identified by histone H3 variant CENPA^2^ and in most seed plants and animals are composed of mega-base length arrays of tandem repeats known as satellites that can phase CENPA nucleosome positions^3–6^. Centromere function is essential across all eukaryotes, yet centromere sequences evolve rapidly, a phenomenon known as the centromere paradox^7^. Comparative centromere sequence analysis between two complete human hydatidiform moles (CHMs) that have fully assembled centromere sequences reveals that only ~63-71% of the sequences can be aligned between the two haplotypes, which highlights the rapid evolution of centromere sequences even within a single species^8^.

Human centromeres are located at α-satellite arrays (α-Sat) comprised of blocks of 171 bp head-to-tail tandemly organized monomers that can differ by 50~80%^9^, but are organized in highly homogeneous higher order repeats (HORs), which themselves have a nested structure^10,11^ in which the most recent and homogeneous HOR that forms the active centromere is surrounded by older more divergent HORs flanked by divergent monomers^10,11^. Based on the layered expansion model of centromeric array evolution, the active α-Sat HOR originates from newly emerged small repeats and expands into a mega-base-sized array within the active centromere while pushing the pre-existing diverged α-Sat HORs to the periphery of the active centromere^9,11^. The copy number (CN) of the active HOR, which indicates the array size of the centromere, varies substantially among individuals (up to ~80-fold)^8^. In addition, the HOR CN between cancer cells and their normal tissue counterparts significantly differ, which reveals that the array sizes can change in the lifetime of an organism^12^. Despite the extreme degree of inter- and intra-individual polymorphism in HOR CNs, the molecular mechanisms that underlie array expansion and contraction, the rate of variation, and consequences of variations are not fully understood.

A widely cited model used to explain the expansion and contraction of satellite arrays involves unequal crossover and gene conversion between sister chromatids during homologous recombination to repair DNA double strand breaks (DSB)^13^. This model, if correct, would predict stochastic expansion and contraction resulting in randomly mutated monomer sequences without any specific structure due to functional constraints^14^. Tandem repeats that are repaired by Single Strand Annealing (SSA), one of several DSB repair pathways, will cause a deletion of satellite repeats^15^. This loss of repeats will eventually shrink centromeric arrays over time unless there is a mechanism that counteracts the loss^6^.

An alternative model is based on break-Induced replication (BIR)^16^ which is a one-sided DSB repair mechanism that can replicate hundreds of kilobases in budding yeast^17^. BIR has been implicated in oncogene-induced DNA replication^18^, replication stress-induced DNA repair synthesis in mitosis (MiDAS)^19,20^, and in significantly elongating the size of the telomere in Alternative Lengthening of Telomere (ALT) positive human cancers^21^. Centromeres are enriched with DNA breaks^22^ that may be caused by replication fork collapse^23^ due to the presence of replication barriers such as the constitutive centromere associated network (CCAN) and non-B form DNA secondary structures^24–29^. Following the 5′-to-3’ resection of a one-sided DSB caused by a collapsed replication fork, the 3′ single-stranded DNA is available for strand invasion, which triggers BIR^17^. Because of the tandemly repetitive structure of satellites, BIR can create deletions, duplications, or neither, depending on the location of re-initiation of a collapsed fork^6^. These outcomes can cause the α-Sat monomer turnover and HOR structure frequently observed in the centromere^14,16,30^. Previous studies at the rDNA repeat arrays in budding yeast have shown that BIR favors out-of-register re-initiation of broken forks leading to array expansion^31^. This expansion bias can counteract the erosion caused by SSA. Frequent dissociation of Polδ from the template DNA and reduced efficiency of mismatch repair during BIR can explain an elevated nucleotide substitution mutation rate in the centromere^17,32^.

These alternative models have not been experimentally validated because of the homogeneity in satellite sequences, their complex organization, and the extremely large size of the centromere. These features of satellite centromeres have hampered centromere experimental biology for decades^6^ until the recent advances in telomere-to-telomere (T2T) assembly^11^.

With the benefit of fully assembled centromere sequences^33^, we measured the CN variation in the chromosome 11 centromeric HOR D11Z1 at intervals of ~20 somatic cell divisions across subclones of the K562 and U2OS cell lines. We found that the D11Z1 CNs vary among subclones of U2OS with a change frequency from ~7% of subclones to ~100%. Using this basal rate of change that we identified, we set out to test by mutation the involvement of the homologous recombination protein RAD52 and helicase PIF1^34,35^. Our data indicates that both RAD52 and PIF1 are required to cause a large change in CN during somatic cell divisions, suggesting that these changes occur via BIR. Our findings provide insight into the mutational mechanism that underlies rapid centromere evolution, while offering a tool to further investigate the consequences of centromeric array variation, which influences the occurrence of genetic disorders, and facilitates speciation^1,7,36^.

## Results and Discussion

### Comparison of ddPCR assays for HOR copy number measurement

The molecular mechanisms that contribute to extensive array length polymorphism during mitosis in mammalian cells are unclear. Recent studies have demonstrated that centromeric array sizes can be experimentally estimated using a droplet digital PCR (ddPCR)-based method^12^. With the recent T2T CHM13 genome assembly^33^, primers can now be designed to target a single, unique amplicon for each HOR based on polymorphisms that differ between monomeric units. Because the α-Sat is composed of repeats of HORs, if the HOR CN per array is known, then the size of an array can be estimated by multiplying the size of the HOR unit by the CN (Figure 1A). For example, if the experimental D11Z1 HOR CN is 3600, then by multiplying the unit size, which is 855bp, the size of the entire D11Z1 array is ~3.07 Mb. The ddPCR is a reference-free quantification method with a 2-fold greater sensitivity than qPCR. This method allows us to quantify the CNs of HORs within a chromosome-specific array by partitioning every copy of the HOR within an α-Sat array that is isolated by restriction enzyme digestion to >18,000 droplets (Figure 1B). The droplets are counted by the machine and used to calculate the CN. The HOR CN is normalized using the CN of a single-copy gene located on the same chromosome as the HOR being measured in a parallel reaction to ensure that the HOR CN reflects a single chromosome array.

**Figure 1.**
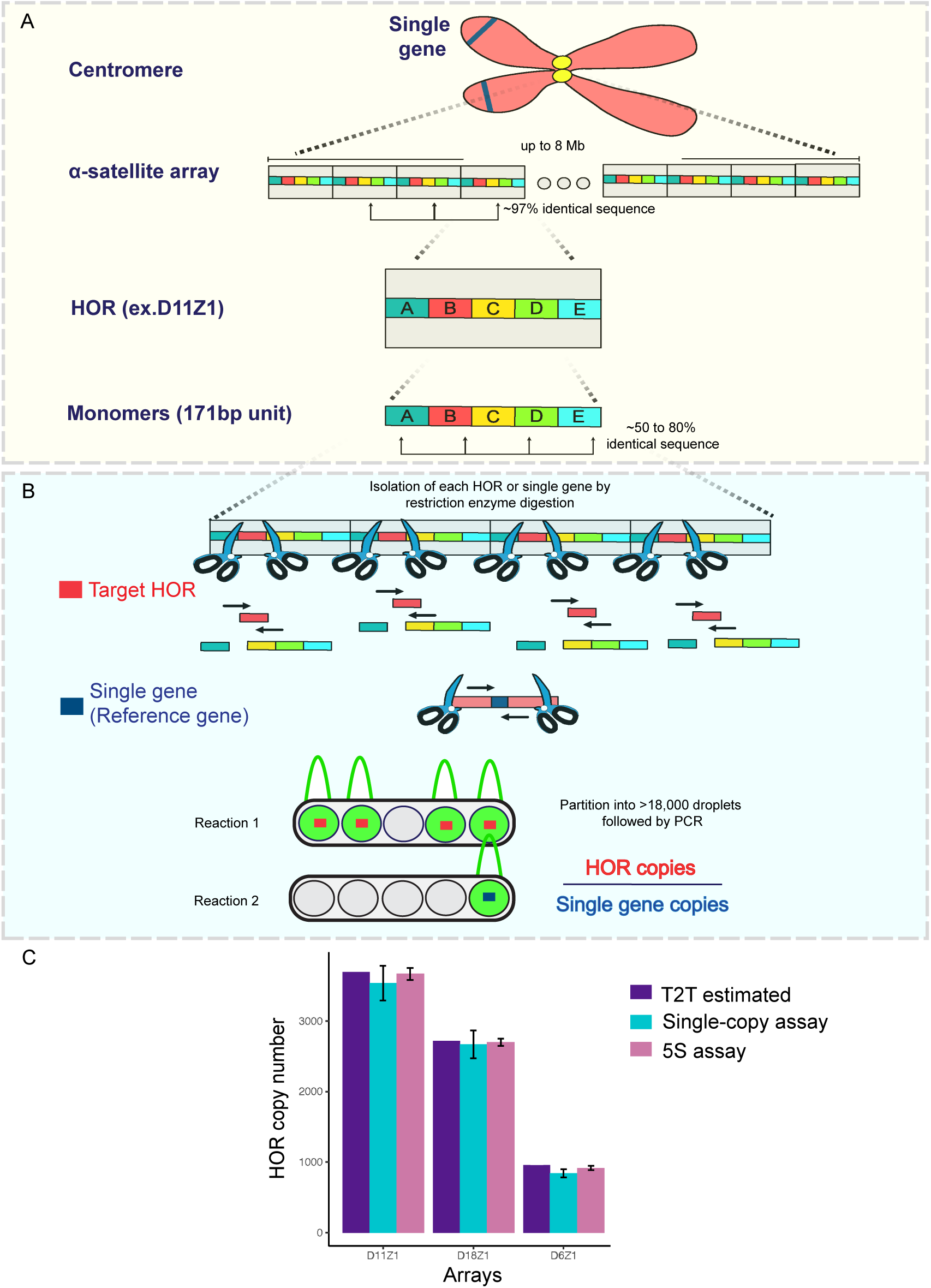
Organization of human centromeres and copy number quantification using ddPCR-based assays. (A) Schematic of the human centromere. Higher-order repeats (HOR, grey box) comprise tandemly oriented 170-171 bp monomers (colored boxes). Specific HOR copy number (CN) can be quantified based on sequence identities between HORs and polymorphisms present in monomers. (B) Schematic of single-copy assay workflow. Each HOR (red box) or single-copy reference gene on the same chromosome (dark blue box) in a subclone is isolated by restriction enzyme digestion, partitioned into >18,000 droplets, and simultaneously amplified using HOR-specific or single-gene primers (black arrows) in separate reactions. The droplets that contain targets (green peaks) are counted by signal amplitude and the CN is calculated. The HOR CN per array is determined by normalization with single-gene copies (e.g. HOR copies/single-gene copies). (C) Histogram showing HOR CNs of D11Z1, D18Z1, and D6Z1 in the CHM13 cell line either measured by the single-copy assay or the 5S assay. Values represent mean ± SD of three independent measurements. For the 5S assay, the CNs of the HOR and 5S were measured and the HOR CN per 5S CN were determined. Next, the 5S and a single gene located on the same chromosome were measured to calculate the 5S CN per chromosome. Finally, this number is multiplied by the HOR CN per 5S CN to calculate the HOR CN per chromosome.

Despite promising applications to centromere biology, the single-copy ddPCR-based assay is associated with an intrinsic error rate between biological replicates of ±10%^12^, probably because the parallel reactions must be carried out at different DNA concentrations. To mitigate this issue, we developed a 5S rDNA probe-based assay that reduces sub-sampling error by measuring both target and reference CNs in the same reaction, using 5S rDNA repeats as the reference gene (Figure S1A). To validate this assay, we measured the D6Z1, D11Z1, and D18Z1 CNs in CHM13 cells using the two methods and compared them with the values that are derived from the CHM13 assembly (Figure 1C)^12^. While the single-copy assay produced values close to those derived from CHM13 assembly, the 5S assay produced values nearly identical to the assembly values with less technical error at the cost of reduced dynamic detection range of the ddPCR.

### Centromeric array CN can change within ~20 somatic cell divisions

A previous study has reported that centromeric array CNs vary substantially between cancers and their counterpart normal tissues, which indicated that centromeric array length alteration can occur in somatic cells^12^. In addition, the pediatric cancers medulloblastoma and acute lymphoblastic leukemia tend to show directionality among the five chromosome specific arrays measured, such as all gain or all loss in HOR CNs, indicating a more coordinated alternation pattern^12^. These coordinated patterns are especially useful to increase the sensitivity of the ddPCR method, since the HOR CNs estimated by ddPCR are an average over homologous chromosomes. This could cause the change in individual chromosome arrays to be masked when the direction of HOR CN change varies between events within the pool of different homologous chromosomes. Therefore, we reasoned that pediatric cancers may be an ideal system because of the coordinated alteration patterns in HORs witnessed in the previous study. We chose the pediatric osteosarcoma cell line U2OS because its telomere maintenance mechanism is known to utilize BIR^21^, which we hypothesized to be the primary mechanism of CN change in centromeres.

We first assessed whether the HOR copy in an array can change during ~20 somatic cell divisions in the U2OS cell line and determined the rate of change that can be measured using the ddPCR-based method. We measured the CNs of D6Z1, D11Z1, D18Z1, and DXZ1 centromeric arrays in single-cell-derived subclones of the U2OS cell line that had undergone ~20 somatic cell divisions, which we named Group1, along with the parental cells that were frozen down right after single cell isolation so that we could identify any subsequent changes that occurred after isolation (Figure 2A). The HOR CN is normalized by the single-gene CN to determine an average HOR CN per array or allele. To identify the subclones that had changed HOR CN significantly since the time of single cell isolation from the parental cells, Tukey’s Honestly Significant Difference (HSD) test was conducted between parental measurements and subclone values following ANOVA test. Three out of seven subclones of Group1 showed an expansion in the D11Z1 CN, two subclones showed a decrease in D18Z1 CN, and one subclone showed an increase in D18Z1 CN (Figure 2B-C). While the frequency of change is the same between D11Z1 and D18Z1, the magnitude of the change was greater in D11Z1 than D18Z1 (30% vs 22% maximum). We repeated the measurements using the 5S assay and obtained essentially identical results, which validated the changes we observed (Figure S1B). Therefore, the magnitude of CN changes occurring in U2OS cells is above the technical error threshold and can be confidently measured using the single-copy assay. While D11Z1 and D18Z1 CNs changed, there were no CN changes in the D6Z1 and DXZ1 subclones (Figure 2D-E).

**Figure 2.**
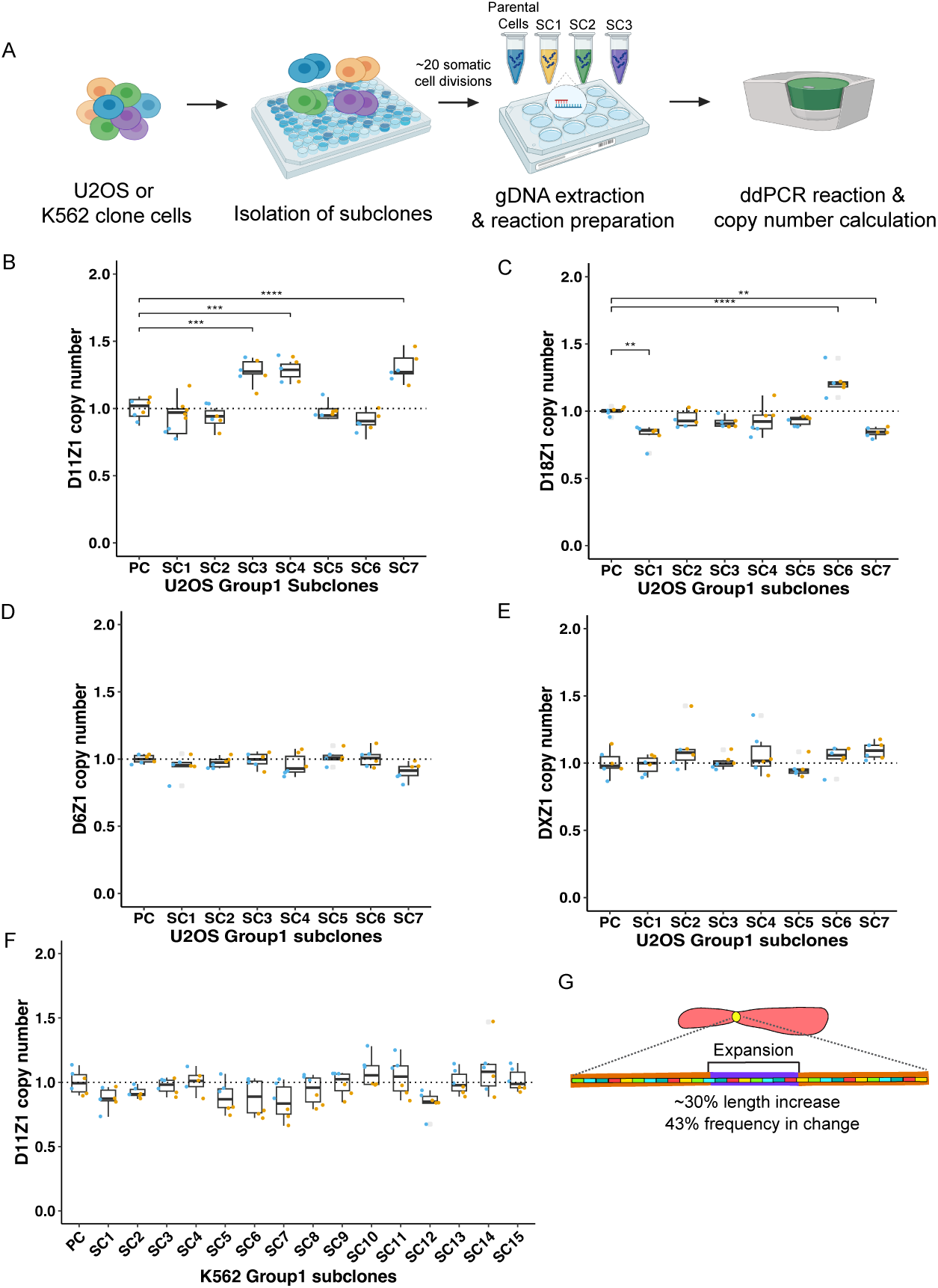
Centromere arrays can expand and contract within ~20 somatic cell divisions. (A) Experimental scheme to quantify HOR CN within somatic cell divisions using single-copy ddPCR-based assay (image created with BioRender.com). (B-E) Box-whisker plots showing the D11Z1, D18Z1, D6Z1, and DXZ1 CNs in U2OS Group1 subclones. In these and subsequent box-whisker plots, each dot indicates a single PCR reaction, which is normalized by the mean of the parental cell (PC) HOR CN (dotted line). Colors indicate technical replicates. Asterisks indicate degree of significance in CN changes between parental cells and subclone pairs determined by Tukey’s HSD test (n=8, Tukey’s HSD, P<0.05). (F) Box-whisker plot showing the D11Z1 CN in K562 Group1 subclones. (n=16, Tukey’s HSD, P>0.05). (G) Cartoon summary of CN changes in U2OS Group1 subclones.

Next, we were curious as to whether the CN changes observed in U2OS cells were broadly characteristic of cancer genomes or were an intrinsic property of the U2OS cell line. Therefore, we also measured the D11Z1 CNs across single-cell-derived subclones of the K562 cell line that had also undergone ~20 somatic cell divisions. Using the single-copy assay, none of the K562 Group1 subclones changed D11Z1 CN (Figure 2F), in contrast to the high frequency and magnitude of change we observed in U2OS cells. However, using the 5S assay, the frequency of K562 clones that changed CN was ~33% (5 out of 15), and all showed expansion, similar to U2OS clones (Figure S1C). The maximum CN alteration observed in K562 was a ~12.4% increase (SC10), which is close to the 10% technical error rate of the single-copy assay, in contrast to the ~30% maximum increase in U2OS CN, likely explaining the failure to detect changes using the single-copy assay in K562 cells. Therefore, we used the single-gene assay in the U2OS cell background for follow-up experiments since it is a more conservative method and the magnitude of change in U2OS subclones far exceeds the technical error threshold.

We conclude that the centromeric array can expand and contract in mitotic cells within ~20 cell divisions in both U2OS and K562 cells, but the magnitude of change is far less in K562. One difference between these cell lines is that U2OS cells undergo BIR-mediated ALT^21^, which is associated with a mutation in the *ATRX* gene that results in a short-lived, truncated protein^37^. ATRX can form a complex with cohesin and MeCP2^38^, and knockout of ATRX causes a defect in cohesion at telomeres, where loss of cohesin between sister chromatids facilitates non-allelic telomere interactions^39^. ATRX depletion likewise causes cohesion defects at centromeres^40^, and we hypothesize that this facilitates non-allelic, out-of-register BIR in tandem centromere arrays, resulting in greater changes in D11Z1 CN in U2OS cells than in K562 cells. Similarly, disruption of cohesion between sisters at the 35S rDNA locus in budding yeast results in the amplification of rDNA through out-of-register replication^41^.

### Expansion occurs more frequently than contraction in D11Z1

The unequal crossing-over model predicts erosion of the centromere over time because any broken replication forks that are repaired by SSA will lead to a deletion of tandem repeats such as HORs^13^. This will inevitably shrink the array unless there is a selective pressure that counteracts the loss so that mega-base array lengths that are seen in most human centromeres are maintained. We had hypothesized that out-of-register re-initiation of replication behind the fork would occur more frequently because the DNA behind the fork is more accessible compared to the positively supercoiled DNA in front of the fork, which would favor expansion^6^. Therefore, we wondered whether centromeric arrays in Group1 that had expanded would continue to expand or would contract.

To this end, we isolated single cell subclones from the two subclones (SC3 and SC4) that increased the D11Z1 CN from the U2OS Group1 (Figure 3A) and allowed them to expand another ~20 somatic cell divisions (Group2). We then measured the D11Z1 CN across Group2 samples and compared them to the corresponding parental HOR CNs. The frequency of CN change in SC3 Group2 was ~42%. At the most extreme CN change, SC3 increased HOR CN ~30% compared to the parental cells of Group1 (Figure 2B) and its subsequent subclone, SC 3.11, gained an additional ~43% in CN resulting in a total expansion of ~86% from parental cells to the SC3.11 cells (Figure 3B). Unexpectedly, half of the Group2 subclones of SC4 decreased the D11Z1 CNs close to the parental cell value of Group1 (Figure 3C). This extreme drop was only observed in these subclones, which were derived from a parental cell that had among the highest starting CN. This led us to examine whether the starting parental CN might influence the direction of change in their subclones. Therefore, we selected SC3.3, which retained a high D11Z1 CN similar to its parent SC3, and SC4.4, which showed a decrease in CN compared to its parent SC4. Single cells were isolated from SC3.3 and SC4.4 (Group3), underwent ~20 somatic cell divisions, and the D11Z1 CNs were measured (Figure 3A). The frequency of CN change for Group3 of SC3.3 was ~7% with only one subclone contracting (Figure 3D). Interestingly, the magnitude of the decrease observed in the SC3.3.6 subclone, whose SC3 parent had a similarly high CN as SC4, was similar to the decrease observed in Group2 of the SC4 subclones (Figure 3A). Finally, all Group3 subclones of SC4.4, which had previously contracted, increased CN (Figure 3E). Among all 55 subclones from U2OS Groups 1-3, ~35% showed expansion and ~13% contraction in D11Z1 CN. This matches our hypothesis that expansion is favored over contraction. While contraction can occur during somatic cell divisions, this only occurred when the subclones were isolated from parent cells with high CN. Therefore, it is tempting to speculate that there might be a homeostasis mechanism that sets an upper limit to centromere CN, analogous to the mechanism that constrains rDNA copy number^31^.

**Figure 3.**
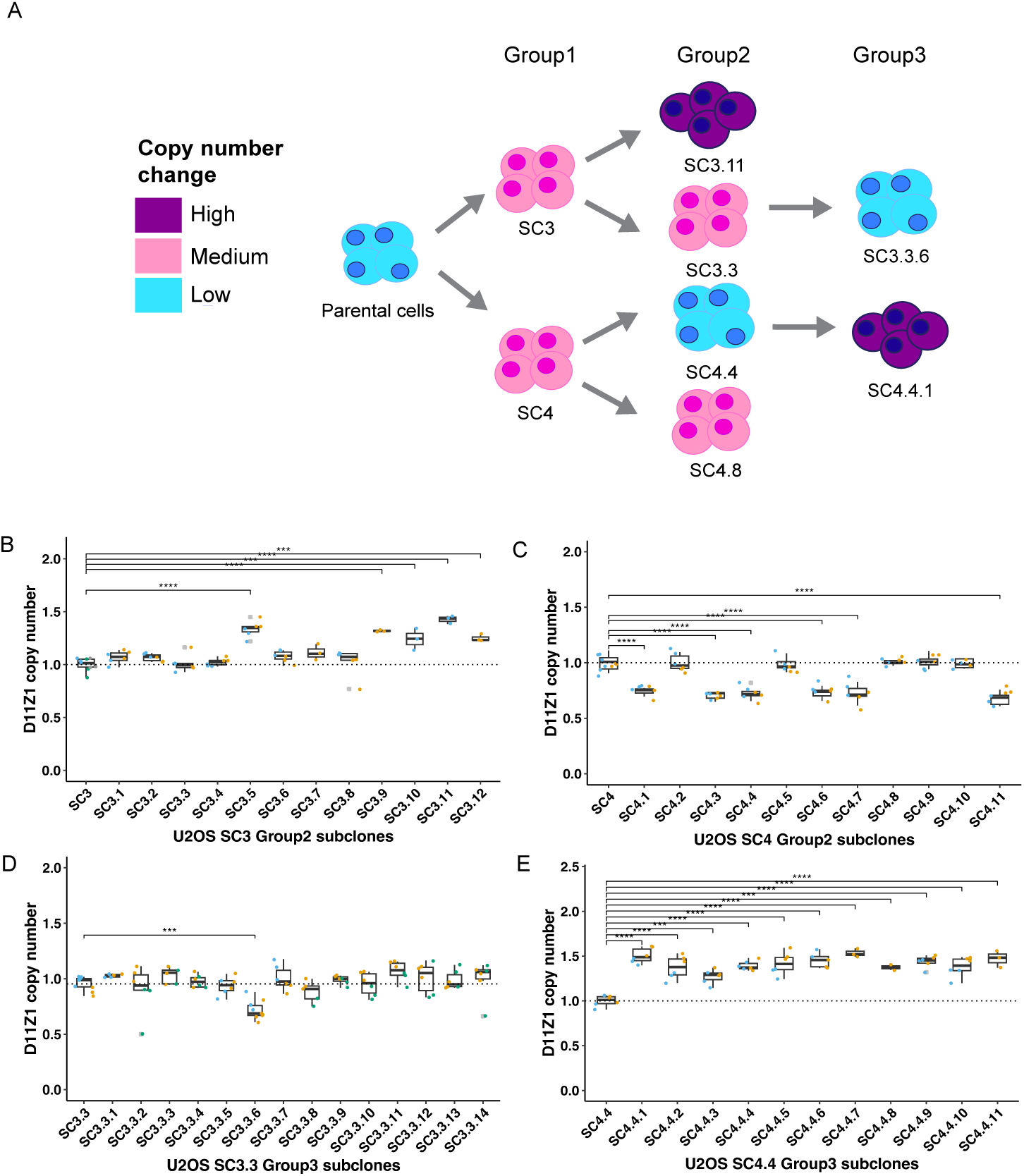
Expansion of centromere arrays is favored over contraction. (A) Schematic of single cell isolation and D11Z1 CN changes over time. Relative magnitudes of D11Z1 CN changes are indicated by colors. (B-E) Box-whisker plots showing D11Z1 CNs in SC3 Group2 (n=13, Tukey’s HSD, P<0.05), SC4 Group2 (n=12, Tukey’s HSD, P<0.05), SC3.3 Group3 (n=15, Tukey’s HSD, P<0.05), and SC4.4 Group3 (n=12, Tukey’s HSD, P<0.05) subclones. Individual subclones are identified as follows: parental cell name followed by a period and subclone number (e.g. SC3.3).

### Centromeric array CN alteration requires both RAD52 and PIF1

BIR-mediated satellite expansion/contraction is a compelling model that can explain unique characteristics of the αSat such as extreme length, complex repeat structure, sequence turnover, and high substitution mutations^16^. During the broken replication fork repair process in yeast, BIR can copy more than 100kb with up to a 1000x higher base substitution mutation rate compared to S-phase DNA synthesis^17^. In addition, out of register re-initiation of replication during BIR repair can cause deletion or addition of multiple copies of repeats resulting in monomer turnover^30^. However, neither the BIR mechanism nor a dependence on the proteins that are required for BIR have been experimentally validated during centromere sequence synthesis.

Thus, we sought to test BIR as the molecular mechanism underlying rapid centromere evolution^6,16^ based on the frequency of D11Z1 array change that we established in our U2OS assays. BIR can occur via the RAD52-dependent pathway, which is well-known for ALT telomere maintenance and MiDAS^21,42^. RAD52-dependent BIR has previously been suggested to mediate centromere expansions in U2OS cells^43^. Mammalian RAD52, which has strand annealing activity, facilitates strand invasion by forming a displacement loop which, in turn, promotes initiation of DNA replication after fork collapse in BIR^19^. PIF1, which is an evolutionarily conserved 5’-to-3’ helicase, is indispensable during BIR for initiation of Polδ DNA synthesis in budding yeast^44^ (Figure 4A). Both RAD52 and PIF1 depletion suppressed BIR-mediated repair of DSBs that were induced by endonuclease I-SceI in U2OS cells^19,35^. In addition, PIF1 is important for BIR-mediated repair of collapsed forks induced by replication stress, resulting in much longer tracts of DNA synthesis than at endonuclease-generated DSBs^35^

**Figure 4.**
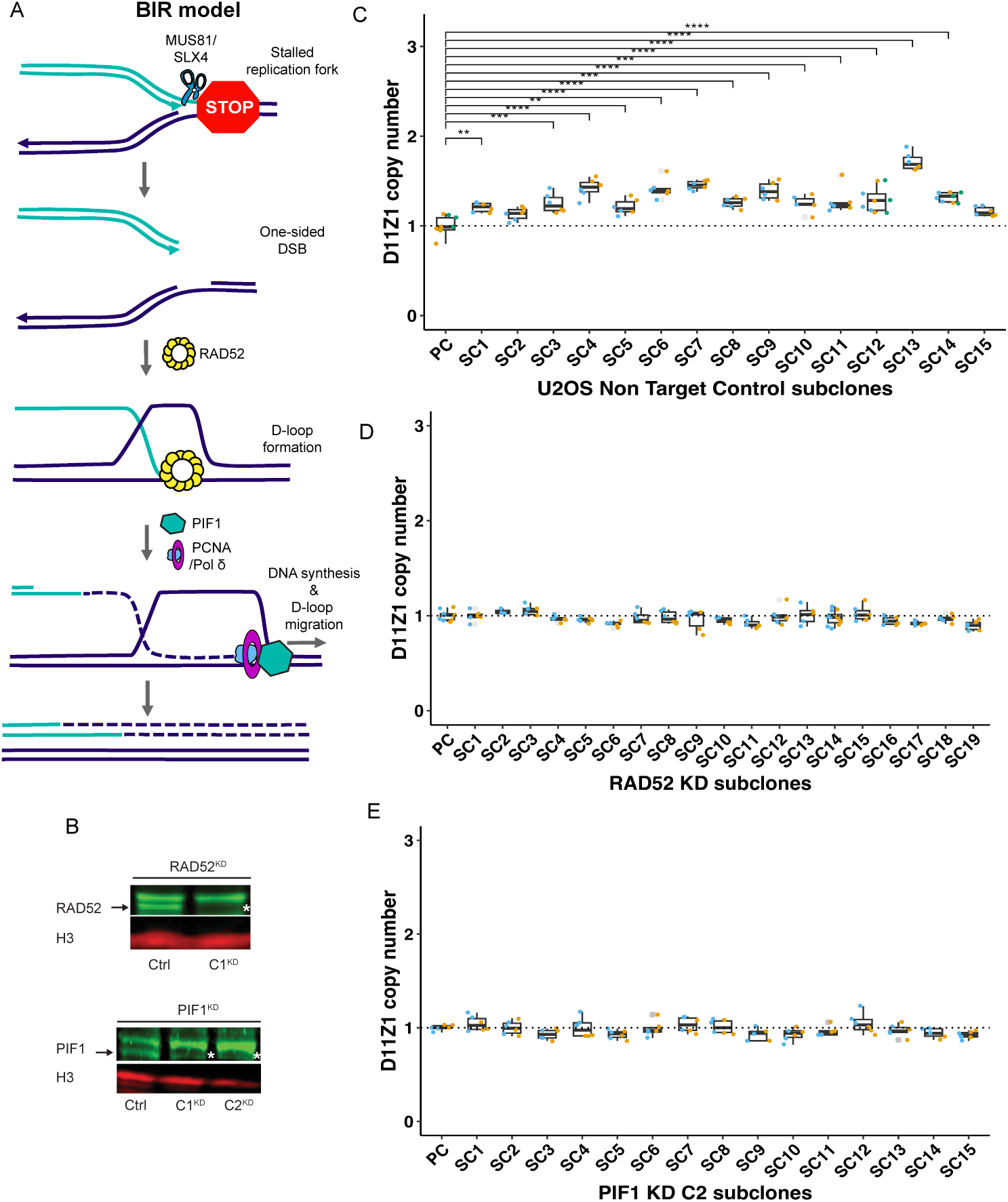
RAD52 and PIF1 are required for D11Z1 CN changes. (A) Overview of the break-induced replication (BIR) model. (B) Expression level of RAD52 (top) and PIF1 (bottom) detected by western blot analysis. Arrows indicate WT bands present in the non-target control. Asterisks indicate KD. The band above the asterisks or arrows are non-specific bands. (C-E) Box-whisker plot showing the D11Z1 CN in either NTC (n=16, Tukey’s HSD, P<0.05), RAD52^KO^ C1 (n=20, Tukey’s HSD, P>0.05), or PIF1 ^KD^ C2 clones (n=16, Tukey’s HSD, P>0.05).

As RAD52 and PIF1 are essential for BIR but not for cell viability we could use knockdowns of these genes to ask whether BIR is required for the CN changes we observed in U2OS cells. Accordingly, we used the CRISPR-Cas9 system to generate knockdown (KD) cell lines, one with disrupted *RAD52* and two with disrupted *PIF1*, along with a non-target control that maintains undisrupted *RAD52* and *PIF1* genes in U2OS cells. To generate the *RAD52^KD^*cell line, we used two single-guide RNAs (sgRNAs) that target exons 3 and 5, which are important for RAD52 oligomerization. For *PIF1^KD^*, we used two sgRNAs that target exons 2 and 6, which are important for helicase activity. We validated one *RAD52^KD^* clone and two *PIF1^KD^* clones using immunoblotting (Figure 4B). We next isolated single cells from *RAD52^KD^ C1*, *PIF1^KD^ C2*, and the non-target control and cultured them through ~20 somatic cell cycles and measured the D11Z1 CN across the subclones. While the frequency of D11Z1 CN change was ~87% (13 out of 15) in the non-target control (Figure 4C), the D11Z1 CN did not change across the *RAD52^KD^*(Figure 4D) and *PIF1^KD^* clones (Figure 4E). These clear results strongly support our hypothesis that BIR is critical for the array size changes that we observed. We repeated the experiment with another knockdown clone, *PIF1^KD^ C1*. The frequency of D11Z1 array change was >80% in the non-target control (Figure S2A), yet no subclone changed D11Z1 CN in the *PIF1^KD^ C1* clone (Figure S2B). Because both *PIF1^KD^s* gave the same result, we can rule out random clonal heterogeneity as a cause of the lack of CN change. Together, these findings demonstrate that RAD52 and PIF1 are required for extensive array size change, leading us to conclude that BIR best explains centromere array size alterations in cancer and likely over evolutionary time scales^16^.

In summary, we have shown that the human α-Sat CN can expand and contract within ~20 somatic cell divisions with a range from ~7% to 100% change in frequency increasing by up to ~86% in CN. These CN alterations favor expansion over contraction and require RAD52 and PIF1, suggesting that BIR underlies centromere sequence evolution in somatic cells. Better understanding of the mechanisms responsible for rapid centromere sequence evolution provides opportunities to study the consequences of divergence in sequence and length and also identifies the players that are involved in this process. Understanding CN change may support the development of therapeutic strategies for cancer, where centromere dysfunction is frequently observed^45^.

## Limitations of the study

While the ddPCR-based assay is an advanced method that allows us to study centromere biology, it has limitations. First, the CNs reported represent the average of the chromosomes in the single-cell-derived population cells. While sufficiently robust to establish CN changes over subsequent clonal generations, a highly heterogeneous population could result in false negatives. Second, the maximum dynamic range of the ddPCR precludes determination of the upper limits of an array size due to increased technical error when the genomic DNA input is lower than the required amount. These false negatives may cause the array change frequency to be underestimated.

## Methods

### Cell culture

All cells were maintained at 37°C and 5% CO2 in T-75 flasks. K562 cells were cultured in suspension in Iscove’s Modified Dulbecco’s Medium (IMDM, Thermo Fisher) with 10% heat inactivated fetal bovine serum (FBS, Cytiva). U2OS cells were cultured in Dulbecco’s Modified Eagle Medium (DMEM, Thermo Fisher) with 10% FBS, Glutamax, 100 units/mL penicillin, and 100 μg/mL streptomycin. CHM13hTERT cells were cultured in basal medium with Amnio Max C100 1X (Thermo Fisher) and the Amnio Max C100 supplement (Thermo Fisher). HEK293T cells were cultured in DMEM with GlutaMAX and 100 U/mL antibiotic-antimycotic (Thermo Fisher).

### Single cell isolation

All parental cells were diluted to place 0.5 cells/well into 96-well plates. First, single cells (Group0) were isolated from the U2OS and K562 population cells and grown until 100% confluence in a 12-well plate (~500,000 cells), which is estimated to be ~20 cell divisions. Subsequently subclones (Group1) were isolated into 96-well plates from a clone in Group0 and underwent ~20 somatic cell divisions. Group2 subclones were isolated from either SC3 and SC4 of Group1 and underwent ~20 cell divisions. Group3 subclones were isolated from either SC3.3 or SC4.4 of Group2 and underwent ~20 somatic cell divisions.

### DNA extraction, quantification, and dilution

Genomic DNA of subclones from a 12-well plate along with 2 million parental cells were extracted using DNeasy Blood & Tissue Kit (QIAGEN) following the manufacturer’s instructions. Genomic DNA samples were quantified using the dsDNA ultra-high sensitivity fluorescent assay (DeNovix) and diluted to 2 ng/μL (single gene CN measurement). Two ng/μL samples were diluted 1:20 to ~ 0.1 ng/μL (HOR and 5S CN measurements). Extracted gDNAs were kept at −20°C.

### Centromeric α-satellite repeats measurement by ddPCR

All primer sequences are listed in Supplementary Table S1A. The four different chromosome-specific HOR array primers (D6Z1, D11Z1, D18Z1, DXZ1) and single gene primers (TBP1, C11orf16, MRO, HPRT1) were used for single-copy assays. For the 5S assay, the same HOR primer sets from the single-copy assay were used for HOR amplification along with a 5S primer set. Two separate probes of different color were used to target the HOR and 5S amplicons. All HOR copy numbers were measured by ddPCR following the manufacturer’s protocol (Bio-Rad).

For the single-copy (EvaGreen) assay, each reaction contained 10 μL of 2X ddPCR EvaGreen Supermix, 0.2 μL of restriction enzyme (Alu I or HaeIII), 1 μL of 2 μM primer mix, 1 μL of 0.1 ng DNA (for HORs) or 2 ng DNA (for single copy genes) and 7.8 μL of nuclease-free water. For the 5S probe assay, each reaction contained 10 μL of 2X ddPCR Supermix for Probes (No dUTP), 0.2 μL of restriction enzyme (Alu I), 1 μL of 20X HEX target primer/probe mix (900 nM /250 nM), 1 μL of 20X FAM target primer/probe mix (900 nM /250 nM), 1 μL of 0.1 ng DNA, and 6.8 μL of nuclease-free water. The reactions were incubated at room temperature for 30 min, emulsified with either EvaGreen or a probe droplet generator oil using an automated droplet generator (Bio-Rad), and then transferred to a 96-well plate. The plate was heat-sealed with foil (Bio-Rad) and then a thermocycling reaction was performed using the following temperature profile, where a 2°C/sec ramp rate was applied to all steps: The EvaGreen assay used a 10 min enzyme activation step at 95°C 40 cycles containing a 30 sec denaturation at 96°C and a 60 sec annealing/extension at 56°C, followed by sequential 5 min signal stabilization at 4°C and 90°C and a hold at 4°C. The 5S probe assay used a 10 min enzyme activation step at 95°C, 40 cycles containing a 30 sec denaturation at 94°C and a 60 sec annealing/extension with 2°C/sec ramp rate at 56°C, followed by 10 min enzyme deactivation at 98°C and held at 4°C. Upon completion of PCR, the 96-well plate was transferred to a QX200 droplet reader (Bio-Rad). For the single-gene assay, QuantaSoft software calculated either the HOR or single gene copies/μL that were used to normalize the HOR CN per chromosome. For the 5S assay, the HOR CN was normalized by the 5S CN automatically by QuantaSoft.

### Generation of CRISPR-Cas9 knockdown cells

Two sgRNA oligonucleotide probes targeting different sites in human *PIF1* and *RAD52* or non-target were cloned into lentiCRISPRv2 puro (Addgene). Plasmids that contain each sgRNA were transfected to HEK293T cells using Lenti-X packaging single shots (Takara) for viral packaging. The Lentivirus was harvested at 48 and 72 h after transfection, combined, and centrifuged. The supernatants were concentrated using a Lenti-X concentrator (Takara) according to the manufacturer’s instructions. Viral titers were calculated using Lenti-X GoStix Plus (Takara) according to the manufacturer’s instructions. U2OS cells were transduced with a lentivirus containing polybrene and selected using 1 μg/mL of puromycin for 3 days followed by single clone isolation. Knockdown efficiency of a protein in single cell-derived clones was measured by western blotting.

### Western blotting

Cells that were harvested from a confluent 6-well plate were lysed in RIPA buffer (25 mM Tris-HCl (pH 7.4), 150 mM NaCl, 0.1% SDS, 0.5% sodium deoxycholate, 1% NP-40) supplemented with cOmplete protease inhibitors (Roche) and incubated on ice. Cells were sonicated for 10 sec at 30% amplitude twice and the supernatant was retained after centrifugation. Proteins were quantified using a Pierce BCA protein assay kit. A SDS buffer containing 5% beta-mercaptoethanol (Bio-Rad) was added to the samples. Samples were heated at 95°C and electrophoresed on a 4-12% of Tris-Glycine gel. The gel was transferred to a nitrocellulose membrane. The membrane was blocked with Superblock blocking buffer (Thermo Fisher) and probed for RAD52 (1:100, Santa Cruz), PIF1(1: 100, Santa Cruz), and histone H3 (1: 1000, Cell Signaling Technology). After secondary antibody incubation (1: 20,000, IRDYE 800 donkey anti-mouse IgG, IRDYE 680 goat anti-rabbit IgG), the membrane was imaged using Odyssey imagers.

### Quantification and statistical analysis

For the single-gene assay, the HOR CN was normalized using the single gene copy number as follows (HOR copies per μL/single gene copies per μL)×20 (dilution factor). For the 5S assay, the HOR copy number was normalized using the 5S copy numbers and then multiplied by 5S CN per chromosome as follows. (HOR copies per μL/5S copies per μL)× (5S copies per μL/single gene copies per μL). The D11Z1 copy numbers of subclones were normalized to the mean of parental values. ANOVA tests were conducted among HOR copy numbers of subclones and then significance in HOR copy number change was determined by a Tukey HSD test that compared a pair of HOR copy numbers of a subclone and the value of its parental cells. The P-value is indicated as *P<0.05, **P<0.01, ***P<0.001, ****P<0.0001. All statistical analyses and graphs were performed within the RStudio which is an integrated development environment for R.

## ACKNOWLEDGEMENT

We thank Dr. E. Hatch for U2OS cells, Dr. S. Tapscott and Mr. S. Bennett for technical help, all Henikoff lab members for valuable comments and discussions.

## AUTHOR CONTRIBUTIONS

S.S., P.T., and S.H. designed the study and experiments. S.S. and Y.X. performed the experiments. S.S. analyzed the data. S.H. and R.O.A. supervised the project. S.S., P.T., and S.H. prepared the manuscript with inputs from all authors. This work was supported by the Howard Hughes Medical Institute, NHGRI R01 HG010492-04, and NIGMS R35 GM150532-01 (to R.O.A).

## INCLUSION AND DIVERSITY

One or more of the authors of this paper self-identifies as an underrepresented ethnic minority in their field of research or within their geographical location.

---

**Supplementary figure 1.**
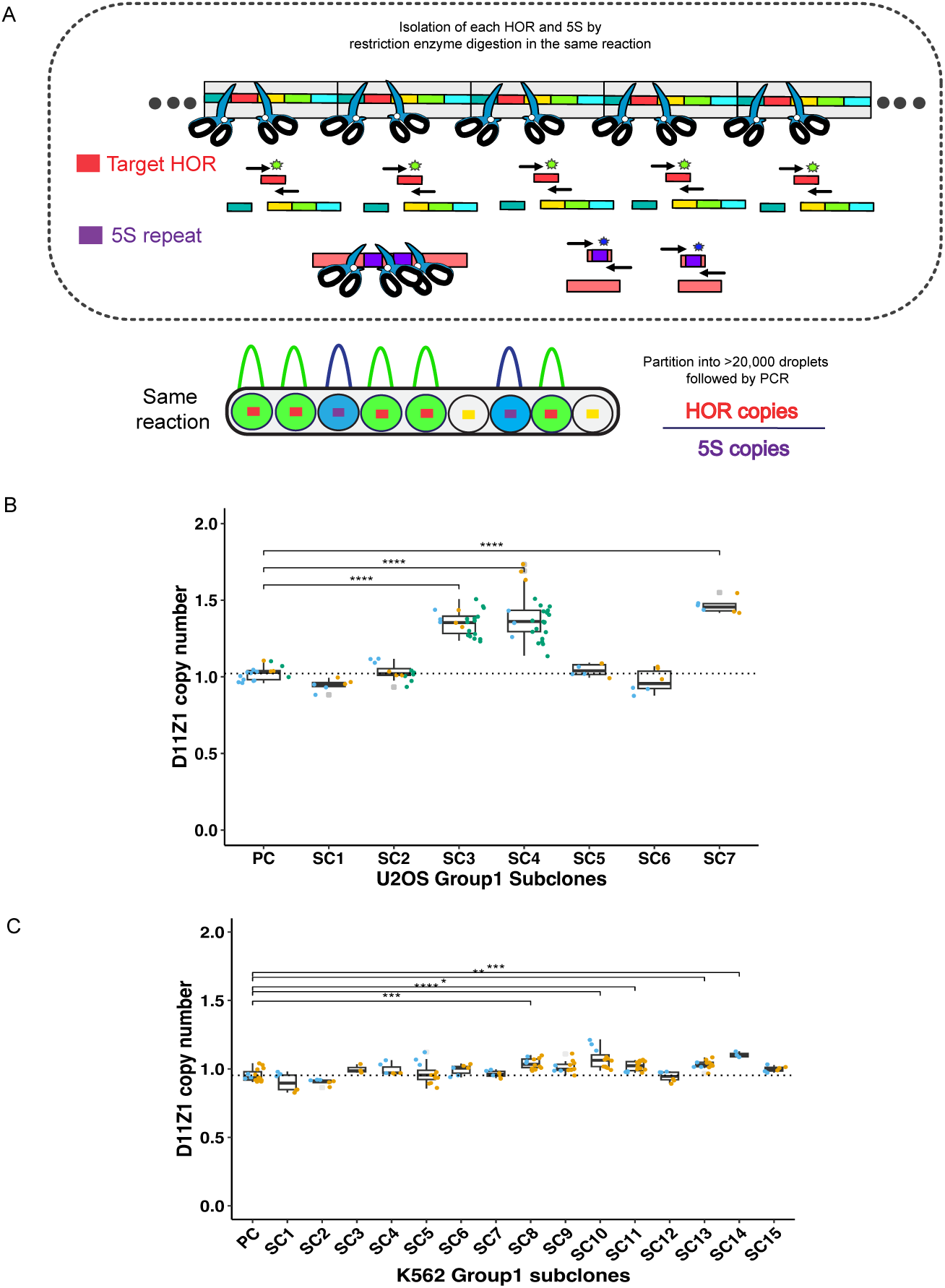
(A) Schematic of 5S assay workflow. Each HOR (red square) and 5S (purple square) from the genome that is isolated by restriction enzyme digestion are partitioned into over 20,000 droplets. Both HOR and 5S targets are simultaneously amplified and bound with the corresponding probes in the same reaction. The droplets that contain any target are counted by two different signal amplitudes and calculated for copies. HOR CN per 5S CN is calculated by Bio-Rad QuantaSoft with 95% confidence intervals. (B) Box-whisker plot showing the D11Z1 CN in U2OS Group1 subclones using the 5S assay. (n=8, Tukey’s HSD, P<0.05) (C) Box-whisker plot showing the D11Z1 CN in K562 Group1 subclones using the 5S assay. (n=16, Tukey’s HSD, P<0.05).

**Supplementary figure 2.**
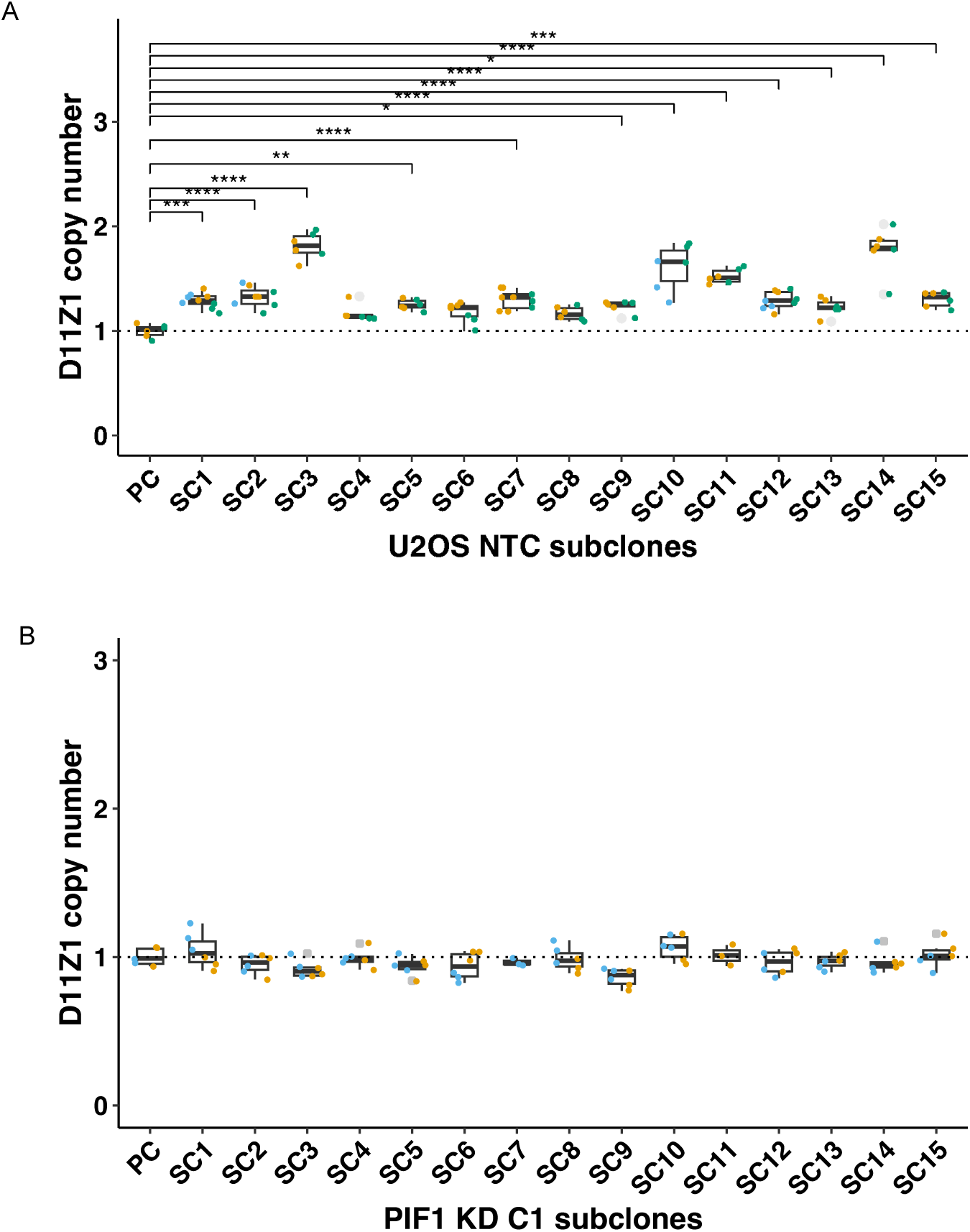
(A-B) Box-whisker plot showing the D11Z1 CN in either NTC subclones (n=16, Tukey’s HSD, P<0.05) or PIF1 ^KD^ C1 subclones (n=16, Tukey’s HSD, P>0.05).

